# Live-bearing cockroach genome reveals convergent evolutionary mechanisms linked to viviparity in insects and beyond

**DOI:** 10.1101/2022.02.03.478960

**Authors:** Bertrand Fouks, Mark C. Harrison, Alina A. Mikhailova, Elisabeth Marchal, Sinead English, Madeleine Carruthers, Emily C. Jennings, Martin Pippel, Geoffrey M. Attardo, Joshua B. Benoit, Erich Bornberg-Bauer, Stephen S. Tobe

## Abstract

Insects provide an unparalleled opportunity to link genomic changes with the rise of novel phenotypes, given tremendous variation in the numerous and complex adaptations displayed across the group. Among these numerous and complex adaptations, live-birth has arisen repeatedly and independently in insects and across the tree of life, suggesting this is one of the most common types of convergent evolution among animals. We sequenced the genome and transcriptome of the Pacific beetle-mimic cockroach, the only truly viviparous cockroach, and performed comparative analyses including two other viviparous insect lineages, the tsetse and aphids, to unravel the genomic basis underlying the transition to viviparity in insects. We identified pathways experiencing adaptive evolution, common in all viviparous insects surveyed, involved in uro-genital remodeling, maternal control of embryo development, tracheal system, and heart development. Our findings suggest the essential role of those pathways for the development of placenta-like structure enabling embryo development and nutrition. Viviparous transition seems also to be accompanied by the duplication of genes involved in eggshell formation. Our findings from the viviparous cockroach and tsetse reveal that genes involved in uterine remodeling are up-regulated and immune genes are down-regulated during the course of pregnancy. These changes may facilitate structural changes to accommodate developing young and protect them from the mothers immune system. Our results denote a convergent evolution of live-bearing in insects and suggest similar adaptive mechanisms occurred in vertebrates, targeting pathways involved in eggshell formation, uro-genital remodeling, enhanced tracheal and heart development, and reduced immunity.

---

Insecta is one of the most diverse animal classes with the highest number of living species, which have colonized most habitats spanning terrestrial, freshwater, and aerial environments (1). Insects have adapted to numerous ecological niches and display a wide range of phenotypic traits. Insect biodiversity is a valuable resource for ecosystems and the source of many new scientific discoveries (1). For instance, insects exhibit a broad spectrum of complex traits such as sociality (solitary, gregarious, sub- to eusociality), metamorphosis (a-, hemi-, pauro- and holometabolous development), and reproductive modes (ovi- to viviparity). While the majority of insects are oviparous (egg laying), viviparity (live birth), both facultative (including ovoviviparity) and obligate, has emerged independently over 65 times across insect evolution (2–4). Among all viviparous insects, the pacific beetle-mimic cockroach, *Diploptera punctata*, and tsetse, stand out by their evolutionary adaptations to have yielded specific organs that house developing progeny and produce protein-rich nutrition, which are funcionally equivalent to placental structures in vertebrate (5, 6).

True viviparity is a reproductive mode in which females harbor developing embryos and other juvenile stages within their reproductive tracts until giving birth to live and active offspring (7). In contrast, oviparity describes the reproductive mode whereby females lay eggs, while embryogenesis as well as other early development stages occur outside the female body (7). Viviparity has gradually evolved from oviparity repeatedly and independently across the tree of life, for instance, within reptiles, mammals, fish and insects (3), suggesting this is one of the most common types of convergent evolution among animals.

Despite broad physiological and morphological differences among viviparous animal clades, the emergence of viviparity has led to similar physiological, morphological, and immunological changes to the female reproductive tract for vertebrate systems (8–10). This transition requires numerous adaptations, observed in both mammalian and reptile lineages, including eggshell reduction, delayed oviposition, enhanced supply of water and nutrition to the embryo by the mother, enhanced gas exchange, and suppression of maternal immune rejection of the embryo (8, 11). The adaptation to viviparity requires acceptance of a developing non-self organism by the mother. In mammals, repression of maternal immunity towards placental cells is essential for successful pregnancy (12), while in reptiles some viviparous squamates display reduced immunocompetence during pregnancy (13). In both mammals and viviparous reptiles, genes and gene families share similar immuno-repression roles in the uterus and placenta (13).

Most of our knowledge on the molecular evolution of viviparity stems from studies in vertebrates (15). The expansion of genomic resources for insects represents an ideal opportunity for investigating more general insights into the emergence of viviparity and comparing distantly related taxa for convergence. By sequencing and assembling the genome of the only known truly viviparous cockroach, *D. punctata*, we investigated the genomic signatures of insect viviparity comparing three origins of insect viviparity, the obligate viviparous cockroach, the obligate viviparous tsetse (*Glossina morsitans*), and two cyclically viviparous aphids (*Acyrthosiphon pisum*, *Rhopalosiphum maidis*). These three systems use strikingly different form of viviparity (5), which are even more divergent then those in other vertebrate systems. Furthermore, we analyzed the transcriptomes of *D. punctata* and multiple *Glossina* species during different pregnancy stages to unravel patterns of specific gene expression before and during pregnancy. We detected genes and pathways under positive selection and also experiencing variation of selection at each of these transitions from oviparity to viviparity. From the results obtained with our comparative genomic and transcriptomic analyses, we selected candidate genes, whose strong effects on pregnancy were validated with RNA interference experiments in *D. punctata*. Our analyses shed light on the biological bases of the emergence of viviparity in insects, which to a large extent mirror convergent viviparous adaptations in vertebrates despite broad physiological and morphological differences.

## Results

### Genome of the viviparous cockroach

We sequenced and assembled the genome of the viviparous cockroach, *Diploptera punctata* (Blaberidae), with a combination of long-(PacBio, 60x) and short-read (Illumina, 45x) sequencing data. We obtained a highly contiguous (contig N_50_: 1.4 Mb) and complete (97.6% of insect BUSCOs) genome assembly of length 3.13 Gb (estimated genome size based on k-mer distribution of Illumina data: 3.07 Gb). This genome size is considerably larger than that of the closest related blattodean species with a sequenced genome, *Blattella germanica* (Ectobiidae, 2.0 Gb) (16), and in fact is closer in size to the more distantly related American cockroach, *Periplaneta americana* (Blattidae, 3.4 Gb) (17). The differences in genome size do not seem to have been aided by variation in transposable element content as all three cockroach genome assemblies exhibit similar proportions of repetitive elements: 54.3% in *D. punctata*, 54.7% in *B. germanica* and 57.8% in *P. americana*. Similarly, we find no evidence for genome size being driven by proteome expansion as we identified 27,939 protein-coding genes similar to the number of proteins first reported for *B. germanica* (29,216), both greater than in *P. americana* (21,336, Fig. 1).

**Fig. 1.**
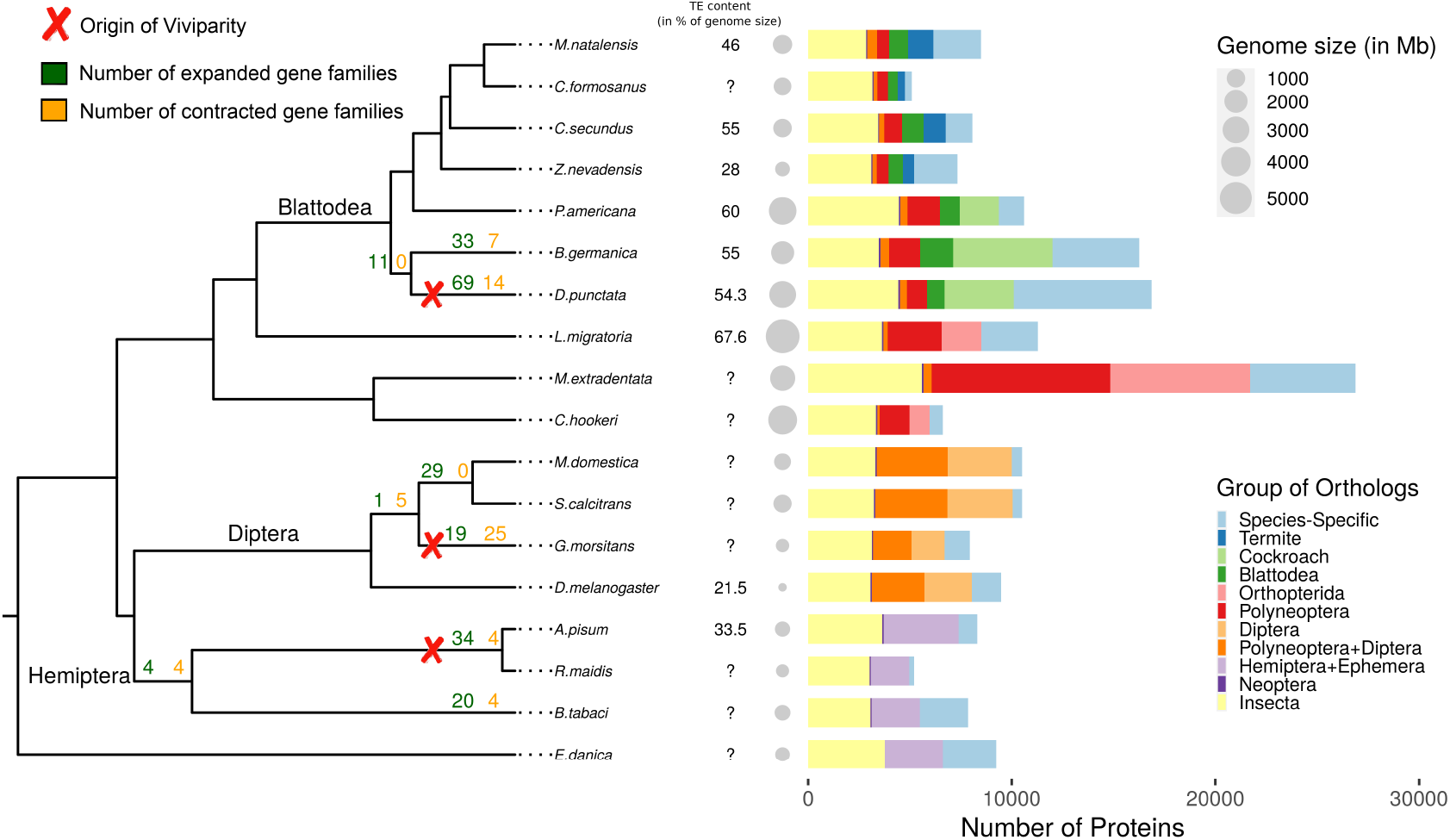
Genomic features associated with viviparity across 3 insect groups. The phylogenetic tree (on the left, made with ggtree (14)) depicts the evolutionary history of 18 insect species along which viviparity originated 3 times independently: in aphids, flies and cockroaches (red cross). Right of the phylogenetic tree, the numbers represent the proportions of transposable elements within each species genome (in %), the grey dots depict their genome size, and the barplot the number of orthologs shared among all 18 insect species, each clade, within Blattodea, within cockroaches, with termites, and those that are species-specific.

### Gene family evolution related to insect viviparity

To understand mechanisms underlying adaptations to viviparity in insects, we compiled a data set comprising genomes and proteomes of 18 insect species from 3 different insect orders (Blattodea, Diptera, and Hemiptera), in which viviparity has arisen independently (Fig. 1). An analysis of gene family size variation from orthogroups sizes defined by OrthoFinder (18) using CAFE (19) across these species revealed several significant gene family expansions and contractions that were shared between independent origins of viviparity. While none of these were shared among all 3 origins of viviparity, 5 expanding gene families were shared between *D. punctata* and *G. morsitans*, 3 between *D. punctata* and aphids and 3 between *G. morsitans* and aphids. Interestingly, most of these gene families experiencing significant expansions across pairs of viviparity origins were either related to protein ubiquitination or chromatin remodelling (Table 1).

**Table 1.** Gene family experiencing either expansion or contraction shared among insect viviparous origins.

| Orthogroup | Reference | Putative function |
| --- | --- | --- |
| EXPANSIONS SHARED BETWEEN *D. PUNCTATA* & *G. MORSITANS* |  |  |
| OG0000835 | *polybromo* | chromatin remodeling |
| OG0001065 | NADH dehydrogenase | oxidative metabolism |
| OG0001123 | NADH dehydrogenase | oxidative metabolism |
| OG0001271 | NADH dehydrogenase | oxidative metabolism |
| OG0001573 | *ref(2)P* | autophagy |
| EXPANSIONS SHARED BETWEEN *D. PUNCTATA* & APHIDS |  |  |
| OG0000078 | *His3* | nucleosome formation |
| OG0000160 | *His1* | nucleosome formation |
| OG0000345 | Poly(ADP-ribose) polymerase catalytic domain | DNA damage & ubiquitination |
| EXPANSIONS SHARED BETWEEN *G. MORSITANS* & APHIDS |  |  |
| OG0000062 | DNA polymerase type B | DNA replication |
| OG0000395 | *Ulp1* | SUMOylation & ubiquitination |
| OG0000630 | Collagen triple helix repeat | tissue structure |
| CONTRACTION SHARED BETWEEN *D. PUNCTATA* & APHIDS |  |  |
| OG0000073 | Reverse transcriptase | transcription |
| CONTRACTIONS SHARED BETWEEN *G. MORSITANS* & AND APHIDS |  |  |
| OG0000010 | *CHKov2* | unknown |
| OG0000017 | *Ser12* | serine-type endopeptidase |
| OG0000033 | *Jon25Bi* | serine hydrolase activity |

Fewer overlaps were found between contracting gene families with no overlap between *D. punctata* and *G. morsitans*, only 1 contracting gene family containing a reverse transcriptase domain was shared between *D. punctata* and aphids, and 3 contracting gene families involved in serine enzymatic pathways between *G. morsitans* and aphids (Table 1). In addition, 2 gene families that were expanded in *D. punctata* were contracted in both *G. morsitans* and aphids. These 2 gene families are part of the serine protease and ecdysteroid kinase-like protein families. Interestingly, serine protease and peptidase inhibitors have been associated with placental growth in lizards (13). The contraction of serine protease gene families in viviparous insects might serve a similar role as inhibitors in reptiles allowing placenta formation.

To link broad functions with gene duplication and loss events during the adaptation to viviparity, we performed enrichment analyses of Gene Ontology (GO) terms among all expanded and contracted gene families, using Fisher’s exact test within topGO package (20) (see Methods). More GO terms were found to be significantly enriched for expanded gene families (*P* < 0.05), with 72, 95, and 142 significant GOs in *D. punctata*, *G. morsitans*, and aphids respectively, compared to GO terms enriched for contracted gene families, with 44, 94, and 4 significant GOs in *D. punctata*, *G. morsitans*, and aphids respectively. The functional categories “negative regulation of chromatin silencing” and “nucleosome assembly/organisation” were found to be significantly enriched for expanded gene families in all origins of viviparity. In addition, functional categories involved in eggshell formation, oxidative metabolism, and vitamin metabolism were enriched for expanded gene families in both the Pacific beetle-mimic cockroach and tsetse (Table S1). Shared functional categories enriched for expanded gene families among *D. punctata* and aphids were mainly related with immunity, while the tsetse and aphids shared GO terms related with neurogenesis, pole cells, and response to hypoxia. However, these GO terms were no longer significant after False Discovery Rate (FDR) correction at 20%, as the 11 significant functional categories after FDR were only found in aphids, mainly involved in transposition and chromosome organization (Table S1).

Detecting common patterns of gene family evolution among phylogenetically distant taxa may prove difficult due to a reduced ability to detect sequence homology. Hence, we carried out a more robust estimate of duplication and loss events by comparing protein domains rather than genes (21, 22). While no protein domains were found to be significantly expanded or contracted among all origins of viviparity, we found two domains to be significantly expanded during two transitions to viviparity. The transcription initiation factor IID, 31kD subunit domain (PF02291) was expanded in *D. punctata* and aphids, while the proton-conducting membrane transporter domain (PF00361) was expanded in *D. punctata* and *G. morsitans*. Expansion of transcription factors during viviparous transitions might have aided changes of gene expression over the course of pregnancy, similar to the transposon-mediated increase of transcription factor binding sites during mammal evolution (23, 24).

We manually annotated three classes of chemoreceptors in *D. punctata*, the odorant (ORs), gustatory (GRs) and ionotropic receptors (IRs), which are known to be highly abundant in two cockroach species (17, 25) and are reduced in *Glossina* sp. (26) compared to oviparous flies. Compared to numbers in both the German and the American cockroach, we found strongly reduced numbers of each of these chemoreceptor classes. We annotated 434 IRs, 69 ORs and 261 GRs, which were 32-55% and 28-46% lower than in *B. germanica* and *P. americana*, respectively. Similarly, the predicted number of cuticle proteins in the genome was lower in *D. punctata* when compared to the German (22% increase) and American (27% increase) cockroaches, which also are reduced in viviparous flies (26) relative to oviparous counterparts.

### Genes related to oogenesis, morphogenesis and developement under positive selection

To identify protein coding genes undergoing positive selection during the adaptation to viviparity, we used the adaptive branch-site random effects likelihood (aBRSEL) method in Hyphy (27, 28) on all 4,671 single-copy orthologs who have passed filtering (see Methods). We found 160 of these orthologs to have signals of positive selection along the three transitions to viviparity: 35 in aphids, 72 in *D. punctata*, and 55 in *G. morsitans* (Table S2). Of these orthologs experiencing positive selection, two were under positive selection in more than one viviparous branch: *Mhc* encoding myosin heavy chain protein involved in muscle contraction was under positive selection in *D. punctata* and *G. morsitans* branches; and *Coq3* encoding a methyltransferase protein involved in wound healing was under positive selection in aphids and *D. punctata* branches. Interestingly, wound healing pathway is linked with viviparous reproduction in aphids (29). After correcting for multiple testing, only 2 single-copy orthogroups were under positive selection with an FDR < 10%: an unknown gene with a dynein heavy chain linker protein domain (ortholog to *CG14651* from *D. melanogaster*) in *G. morsitans* and the gene (*Cht11*) encoding a chitinase in *D. punctata* involved in chitin metabolism.

To identify whether positive selection among viviparous origins quantitatively relates to particular functions, we classified orthogroups based on their GO term annotations from *D. melanogaster* orthologs and protein domains from a pfam library (see Methods). Using SUMSTAT (31) with the topGO R package (20) to test for gene set enrichment, we identified (*P* < 0.05), 59, 64, and 104 functional categories that were enriched among positively selected genes in the viviparous cockroach, the tsetse, and the aphid branches, respectively (Fig. 2 A-C). While no specific, enriched GO terms were shared among the 3 viviparous origins, all the branches, where viviparity has arisen, had functional categories enriched among genes under positive selection linked with embryogenesis and organ development (Fig. 2 A-C). Furthermore, all branches of viviparous origins shared positively evolving functional categories related to the alleviation of multiple stressors, cell fate and metabolism (Fig. 2 A-C). We also detected signals of positive selection on genes associated with oogenesis and chitin metabolism in both *D. punctata* and *G. morsitans* (Fig. 2A & B). After FDR correction three functional categories remained significant (FDR < 2 0%) a mong g enes u nder p ositive selection on the aphid branch: “mitochondrial electron transport, NADH to ubiquinone”, “vascular endothelial growth factor receptor signaling pathway”, and “imaginal disc-derived wing margin morphogenesis” (Table S3).

**Fig. 2.**
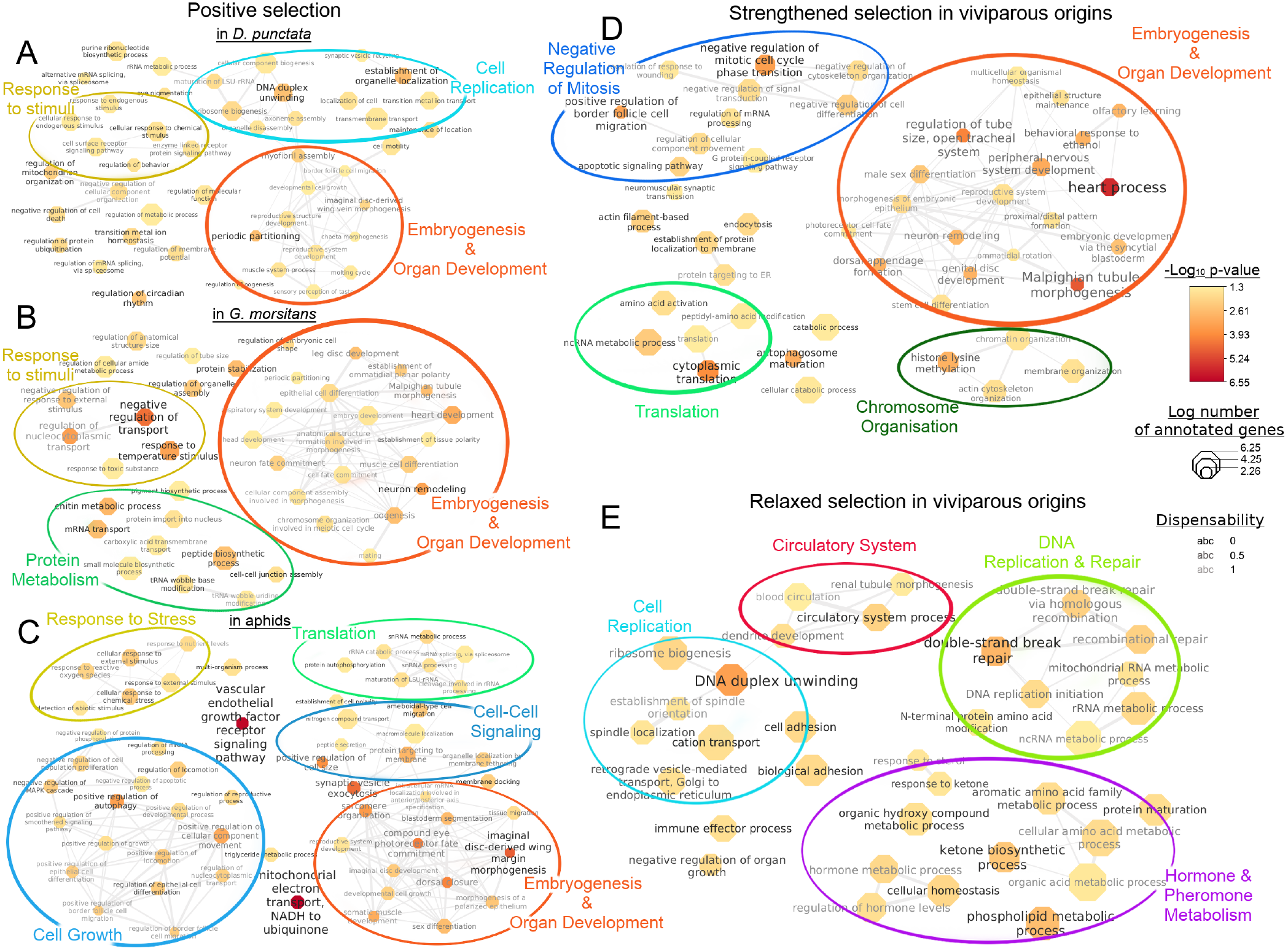
Functional categories involved in the emergence of viviparity in insects. A) Enriched functional categories in genes under positive selection in the viviparous cockroach, *D. punctata*. B) Enriched functional categories in genes under positive selection in the tsetse, *G. morsitans*. C) Enriched functional categories in genes under positive selection in the aphid branch. D) Functional categories enriched for genes under strengthened selection pressure in all branches where viviparity emerged. E) Functional categories enriched for genes under relaxed selection pressure in all branches where viviparity emerged. In each case enriched GO-terms are shown with a p-value < 0.05 and are clustered into broader functions whenever possible, using REVIGO (30) and Cytoscape for visualization.

### Genes with increased and relaxed selection along transitions to viviparity

With the transition to viviparity, it is predicted that there will be stronger selection for adaptations that facilitate live-bearing and relaxed selection on traits associated with evolutionary conflict o ver r esource a llocation. To test this prediction, we identified genes experiencing variation of selection during the transition to viviparity, using the RELAX method in Hyphy (27, 32). This assigns a value determining relaxation or strengthening of selection pressure as well as estimates its significance using m aximum l ikelihood methods, on 4,671 single-copy orthogroups. We identified 2 15 and 112 orthogroups to be under relaxed and strengthened selection, respectively, in the viviparous branches at 10% FDR. From those lists, the highest significant orthogroups (N = 15) experiencing strengthened selection are mostly involved in development, protein metabolism and the excretory system, while those under relaxed selection (N = 11) are mainly involved in cell-cell signalling and cell metabolism (Table S4).

To identify broader functions under differential selection pressure during the transition of viviparity, we tested for GO-term enrichment among genes under relaxed and strengthened selection separately. Only the functional category “heart process” was found to be enriched for genes under stronger selection at 20% FDR. However, using an uncorrected p-value < 0.05 the functional categories enriched among genes under strengthened selection can be grouped into 4 main categories, namely: embryogenesis and organ development; negative regulation of mitotic cell cycle; chromosome organisation; and translation (Fig. 2).

Furthermore, several functional categories under strengthened selection pressure in viviparous species seem to be linked with cell-cell adhesion and signalling (i.e. “establishment of protein localization to membrane”, “endocytosis” and “autophagosome maturation”) and the regulation of gene expression (i.e. “non-coding RNA process”). While functional categories enriched among genes under relaxed selection (*P* < 0.05) do not cluster as well, they can still be grouped within 4 main functions: DNA replication and repair; hormone and pheromone metabolism; cell replication; and circulatory system. In addition to these broad functions, the immune effector pathway is under relaxed selection pressure in branches of live-bearing origin (Table S5).

### Expression patterns related to pregnancy are similar among remotely related viviparous insects

To determine if transcriptional patterns related to pregnancy were shared among independent origins of viviparity, we compared the expression of single copy orthologs between six tsetse species (*G. morsitans*, *G. palpalis*, *G. fuscipens*, *G. austeni*, *G. pallidipes*, *G. brevipalpis*) (26) and *D. punctata* (6). Specifically, the expression in males, early pregnancy, and late pregnancy were compared using a gene co-expression analysis (Fig. 3). These analyses identified that there were specific groups of genes associated with early and late pregnancy in both insect groups (Fig. 3). Early pregnancy showed an enrichment for cuticle changes and associated processes, suggesting that structural changes are a key factor early in insect viviparity (Fig. 3B). The genes associated with late pregnancy are linked with energy and protein production, which highlights the necessary increase in factors to provide nourishment for developing progeny (Fig. 3). Lastly, closer examination of expression of immune-related genes in *D. punctata*, both across pregnancy and in comparison to males and non-pregnant females, revealed immune changes associated with pregnancy (Fig. 3D). These transcriptional shifts associated with pregnancy show functional similarities to those in vertebrate systems (8, 9).

**Fig. 3.**
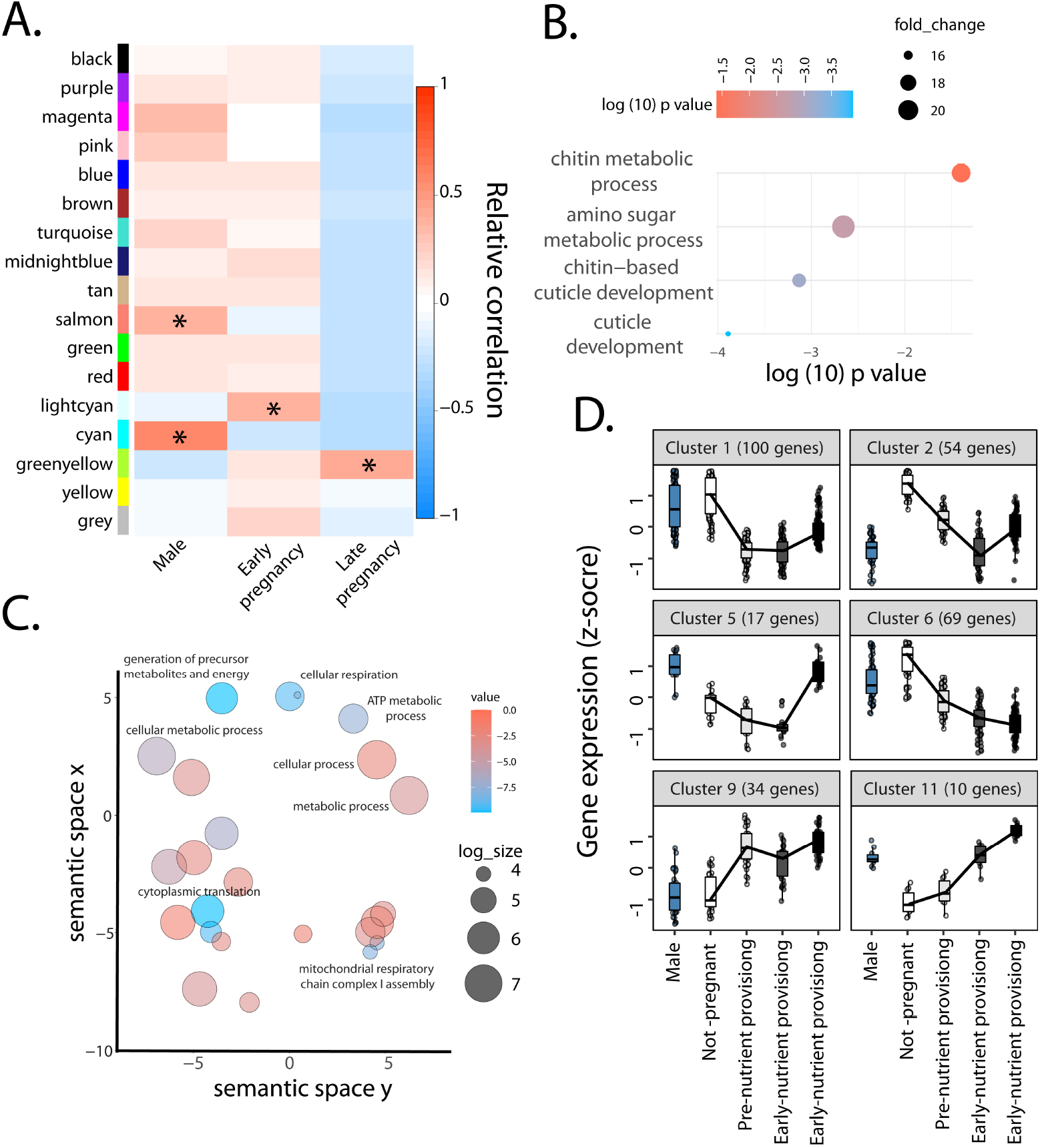
Transcriptional changes associated with single copy orthologs during the course of pregnancy in Pacific beetle mimic cockroaches, *Diploptera punctata*, and tsetse, *Glossina* spp. A) WGCNA-based assessment examining males, early pregnancy, and late pregnancy for *Glossina* and *D. punctata* B) Gene ontology (GO) assessment of modules associated with early pregnancy in *Glossina* and *D. punctata*. C) GO categories associated with late pregnancy in *Glossina* and *D. punctata*. D). Transcriptional changes associated with immune categories over the course of pregnancy in *Diploptera punctata*. Similar immune-pregnancy interactions have been observed in *Glossina* during pregnancy (26). RNA-seq datasets were acquired from (6) for *D. punctata* and (26) for *Glossina* spp.

### RNA interference and immune studies confirm the critical role of viviparity-associated genes

To confirm that specific genes identified by our earlier selection and transcriptome analyses are involved in the process of live birth, we performed RNA interference (RNAi) in *D. punctata* to suppress transcript levels based on previously developed methods (6). These results confirmed that reduction in transcript levels of targeted genes potentially associated with viviparity could increase the rate of abortion and extend the duration of the pregnancy cycle when compared to controls (Fig. 4). These specific genes are involved in protein synthesis and structural aspects, and would not directly be identified as critical to viviparity without our selection and transcriptome analyses. Of interest is the suppression of single milk gland protein, a key component of nutrients for the developing embryo, did not impact pregnancy, likely due to the presence of over 20 similar genes that can compensate to feed the embryos (6, 26) Lastly, immune function was assessed by injection of a bacterium, *Pseudomonas aeruginosa*, which revealed that pregnant females died more rapidly following infection compared to non-pregnant ones. These studies confirm that genomic and transcriptomic factors identified by our analyses are directly linked to cockroach viviparity.

**Fig. 4.**
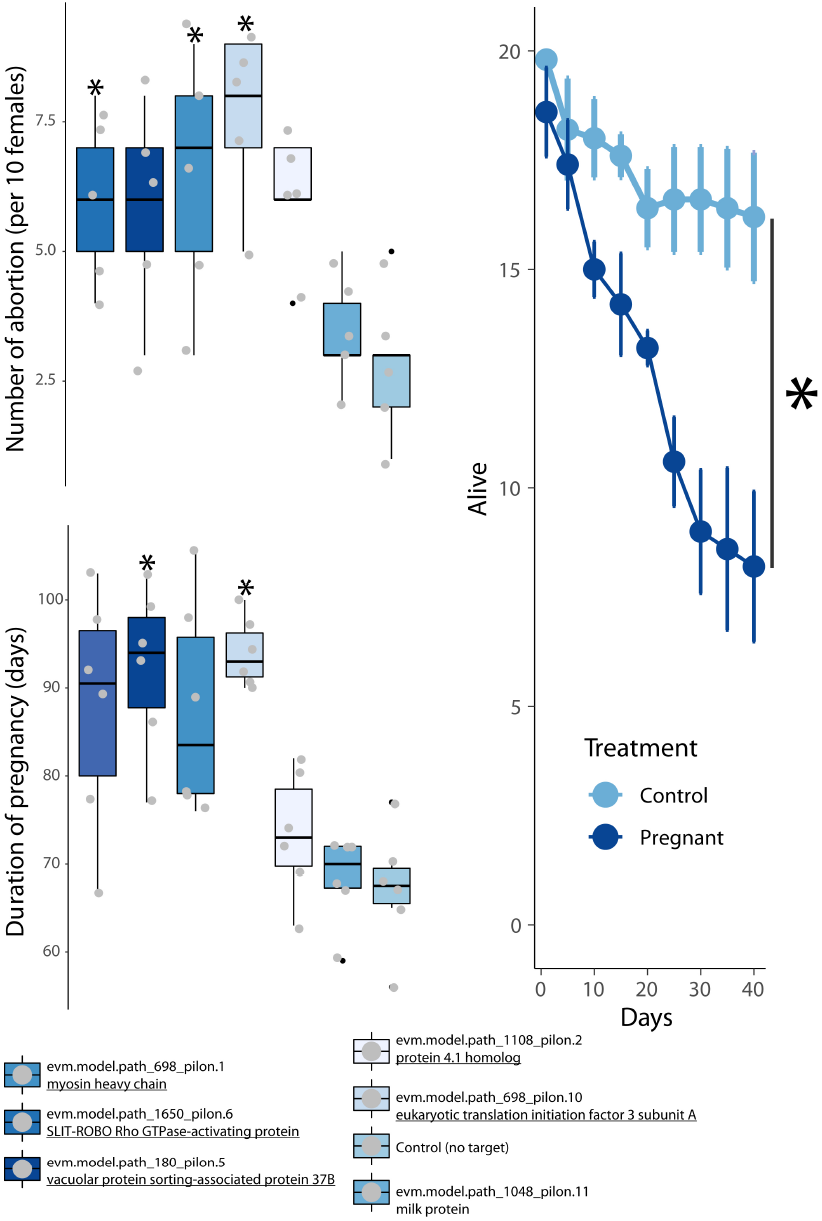
Functional validation of genes and pathways involved in the adaptation to livebearing in the viviparous cockroach. A) RNA interference of genes of interest, selected based on our genomic and transcriptomic analyses, at each of the different pregnancy stages in *D. punctata*, confirmed the role of specific genes during pregnancy leading to disruption of pregnancy and/or delays in birth. B) Immune challenge in pregnant compared to non-pregnant females confirmed the reduced immunity during the course of pregnancy associated with a reduction of survival rate.

## Discussion

Increasing the number of sequenced insect genomes represents a major step towards improving our understanding of the molecular basis underlying adaptive radiation. Comparative genomics of such a diverse animal class provide insights into the key genomic changes along the evolution of insects and also sheds light on the mechanisms by which certain genes and pathways enable the emergence of specific phenotypes. Our genome-wide analyses reveal that convergent adaptations to viviparity in insects are driven by strong positive selection on specific pathways and functional categories, as well as the regulation of specific gene expression patterns during the different stages of pregnancy. Most intriguingly, our results parallel vertebrate adaptations to viviparity with strengthened selection targeting embryogenesis, reproductive system development, tracheal system, and heart development, as well as gene expression patterns during pregnancy linked with reduced immunity and uterine remodelling (Fig. 5).

**Fig. 5.**
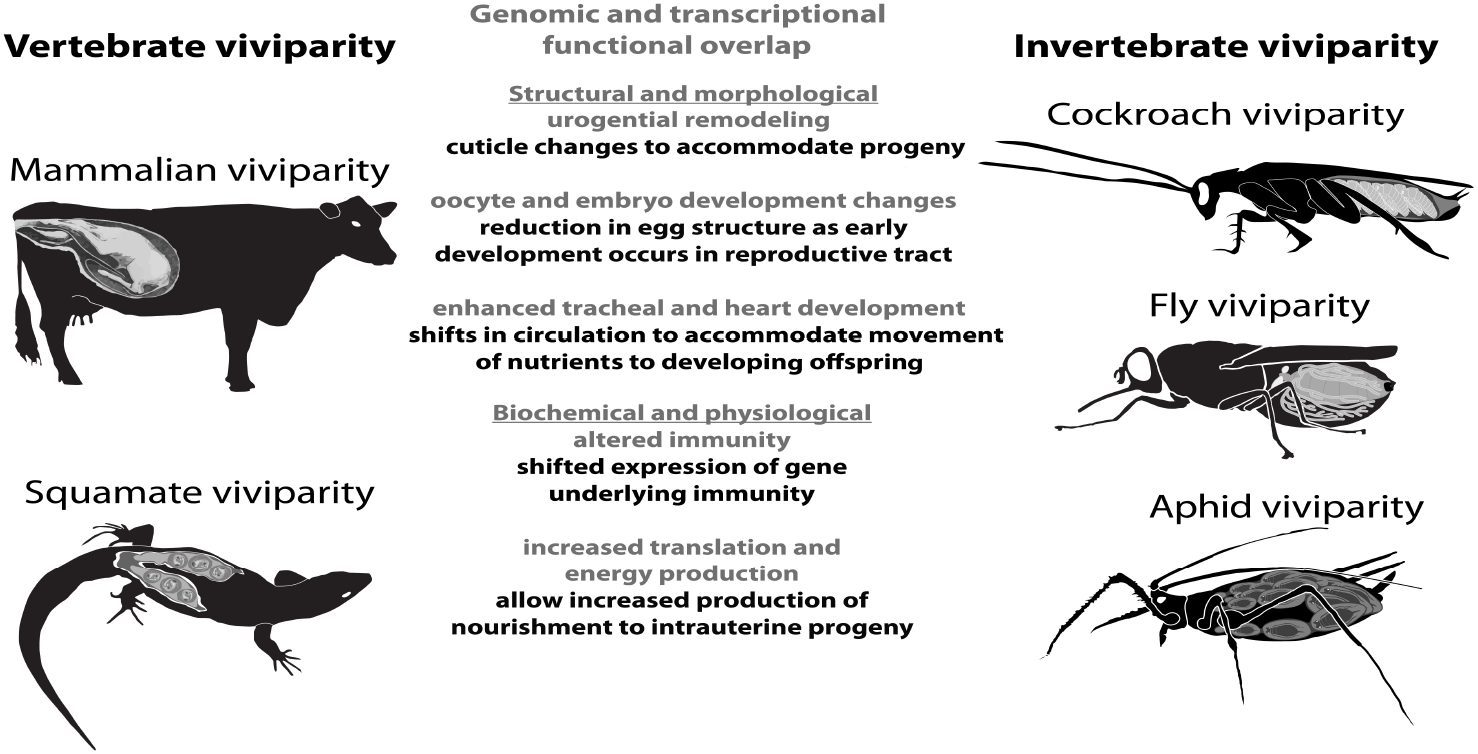
Summary of factors that overlap between vertebrate and invertebrate viviparity. The specific aspects identified were established by a combination of genomic and transcriptomic analyses. Gray indicates common factor between invertebrate and vertebrate viviparity and black is the specific aspect identified in this study for insect systems. Aspects associated with vertebrate viviparity are based upon previous studies (8, 9, 37)

### Morphological and physiological adaptation to viviparity

Our study of gene family evolution revealed a duplication of genes involved in eggshell formation in both the viviparous cockroach and the tsetse, which may have favored viviparous transitions similar to eggshell reduction observed in viviparous reptiles (8, 9). Our genome-wide selection analyses reveal that the adaptation to viviparity in insects is linked with strong positive selection of genes involved in oogenesis in both cockroaches and flies. The transition from oviparity to viviparity is accompanied by a reduction of offspring production per cycle (5). Therefore, the regulation of egg production to reduce offspring numbers might have been attained through amino-acid changes in proteins involved in the development of oocytes. Moreover, in tsetse, alternate oocyte production between left and right ovaries is associated with viviparity, as a consequence of resource constraints (5). In addition to oogenesis and its regulation, our results, for the first time, highlight the rapid evolution of embryogenesis and early development along with the urogenital system development in insects during livebearing adaptation. These are key adaptations to viviparity, which enable the uterus to host the developing embryo (11, 15). The morphological changes of the uterus associated with livebearing is reflected in our results with rapid evolution of genes involved in reproductive system development in all viviparous branches and gene expression changes during pregnancy of both cockroach and tsetse associated with chitin metabolism. In addition, the strengthened selection pressure of genes involved in cell-cell adhesion and membrane trafficking we observed in viviparous branches is likely linked with the emergence of the pseudo-placenta or a placenta-like structure adapted in viviparous insects, which enables enhanced gas exchange and nutrient supply (4, 33, 34). Our results suggest that the rapid evolution of genes linked to the development of the tracheal system and heart underlies the adaptation to enhanced maternal-fetal gas exchange during viviparity, mirroring the adaptation of enhanced angiogenesis during livebearing transition in vertebrates (9). Enhanced gas exchange in viviparous insects might have also been overcome by duplications of genes involved in oxidative or hypoxia metabolism. Furthermore, our results reveal that the emergence of placenta-like structures in insects is associated with enhanced maternal control over fetal development, as genes involved in the embryonic development via the syncitial blastoderm are evolving under strengthened selection pressure along the viviparous branches.

### Immunity and maternal tolerance of the embryo

Another major change linked with live-bearing adaptation in vertebrates is a reduced immunity necessary for not rejecting the developing embryo. In support, during *D. punctata* pregnancy, we observed differential regulation of immune genes especially during early pregnancy, in line with findings in viviparous vertebrates (12, 13). Furthermore, we find immune effectors to be under relaxed selection in viviparous branches, while in many insects immune pathways are expected to be the target of strong positive selection (35, 36), denoting the importance of reduced immune efficiency during live-bearing transition to minimize embryo rejection. Reduced immunity during pregnancy is not only inferred from our transcriptomic and genomic analyses but also revealed by a reduced survival rate of pregnant females after an immune challenge. This difference in survival could also be due to nutritional trade-offs, with the high energetic demands of provisioning developing progeny resulting in reduced immune function, whereas such trade-offs would be less pronounced in oviparous counterparts. The tolerance of the developing embryo within a female’s body might have led to the strong positive selection on pathways related to stress response, which we inferred for all three viviparous insect lineages. Indeed, in all viviparous insect branches, pathways involved in response to chemicals or toxic substances were enriched among positively selected genes, which might indicate adaptations to sustaining embryo growth and development.

### Molecular basis of evolutionary conflict over resource allocations

While profound changes are needed to enable the transition to viviparity as seen above, evolutionary conflicts over resource allocation arise among females against males and offspring (3). Such evolutionary conflicts should result in divergent evolution which could produce similar signals to relaxed selection. One of the major pathways enriched for genes evolving under relaxed selection comprises hormone and pheromone metabolism. In the viviparous cockroach, as well as in tsetse, developing embryos are directly fed with specific nutrients (5, 6). The production of these nutrients is governed by the regulation of Juvenile Hormone (38). Considering the evolutionary conflict over resource allocations, the regulation of nutritional secretions should be the main source of conflict as it represents the main source of nutrients for the embryo. The relaxation of selection on these pathways in viviparous branches could highlight such evolutionary conflict, with balancing selection between females minimizing the production of pregnancy-associated secretions while male and offspring maximize its intake.

### Universal pathways to viviparity

Overall, our study reveals that the viviparity transition in insects is associated with strong positive selection or strengthened selection pressure of genes involved in oogenesis, embryogenesis, tracheal system, and heart development. In addition, our analyses highlight uterus remodelling associated with viviparity in insects detected in a change of gene expression related to cuticle metabolism, as well as a strong positive selection pressure on the urogenital development. Along with pathways involved in uterus remodelling, we found the development of placentallike structures in viviparous insects to be associated with strengthened selection pressure on genes involved in maternal control over embryo development. The viviparous cockroach displays reduced immunity during pregnancy with a reduction of immune gene expression during early pregnancy, and the evolutionary transitions to viviparity seem to have led to the relaxation of selection on immune effectors in all three studied viviparous insect branches. Hormonal changes were also noted in the genomic analyses of invertebrate viviparity, which could be as critical as the hormone shifts and changes that are necessary for vertebrate viviparity (8–10).

Moreover, we found that in all viviparous insect branches non-coding RNA processes, involved in the regulation of gene expression (39), are under strengthened selection pressure. Likewise, the basal evolution of eutherian mammals is associated with bursts of regulatory miRNAs regulating the expression of genes involved in placentation (40), non-coding RNA pathways in viviparous insects might therefore play a similar role. In addition, the formation of the placental tissue in mammals is associated with chromosomal maintenance pathways (41), which could explain the strengthened selection pressure of genes involved in chromosomal organisation in viviparous insect branches. Despite similar pathways shared among the three origins of live-bearing in insects, little overlap was found among genes involved in theses adaptations among the different viviparous insect species. Our results highlight that the transition to viviparity involves similar pathways in both insects and vertebrates, but not necessarily common functional genes (9).

Strikingly, despite broad physiological and morphological differences, the adaptation to live-bearing seems to be universal among animals with pathways. In both arthropods and vertebrates, pathways related to eggshell formation, urogenital remodelling, maternal control of embryo development, tracheal and heart development corresponding to angiogenesis in vertebrates, reduced immunity have undergone fast evolutionary changes and gene expression changes during the course of pregnancy (Fig. 5). Even more remarkably, not only similar pathways but similar evolutionary mechanisms underlie the transition to viviparity in animals, with fast evolution, co-option, gene duplication, and expression changes during pregnancy of genes involved in corresponding functions across different animal taxa (15). Different forms of viviparity as well as ovoviviparity, which can be considered intermediate along the transition from oviparity to viviparity, are well represented in insects (2, 42). Comparative genomics of insects, therefore, represents a great avenue to study in depth the genomic basis of the gradual emergence of live-bearing reproduction mode.

## Materials and Methods

### Specimen Collection

Cockroaches were acquired from the Ohio State University Insectary and maintained according to Jennings et al. (6). DNA was extracted from testes and sequenced at the Centre d’expertise et de services Génome Québec.

### Genome Sequencing and Assembly

The genome was sequenced with a combination of long- and short-read technologies. Using Illumina HiSeq, we generated 147Gb of 150bp paired-end reads (486.8M read pairs), with 500bp fragment size. These reads were quality and adapter trimmed with Trimmomatic (v0.38) (43), resulting in 466.4M read pairs and 136.5Gb. We used these trimmed Illumina reads to estimate the genome size by first calculating kmer-frequencies with Jellyfish (v2.3.0) (44), with a kmer size of 21 and a hash size of 109. The resulting histogram of kmer distribution was then used to model genome size with GenomeScope 2.0 (45), which was predicted at 3.07Gb, with an estimated heterozygosity level of 0.4% and repetitive content of 64.2%.

With 38 SMRT cells on a PacBio Sequel system, we generated 15.4M reads and a total of 164.5Gb of subread sequence data (mean read length: 10 712bp). The PacBio sequences were assembled with MARVEL (46). A database was created using blocksize 250. Then to reduce run times, prior to the first alignment step of MARVEL (daligner), raw reads were masked for repeat regions. This was first carried out only on diagonal blocks (e.g. DB.1 vs DB.1, DB.2 vs DB.2 etc.), then subsequently on a broader diagonal of ten blocks, setting the coverage threshold at 10 and 15, respectively. MARVEL was then run with standard settings on these patched reads. The resulting assembly was polished with the patched PacBio reads that were produced within the MARVEL assembly. For this reads were first aligned against the assembly using nucmer from the MUMmer suite (v4.0.0beta2) (47), then a consensus was created with racon (48). This improved assembly was further polished using the Illumina reads, which were first mapped to the assembly with bowtie2 (v2.3.4.3) (49). The resulting bam file was then used to polish the assembly using Pilon (v1.23) (50). Finally, we removed duplicate contigs with Pseudohaploid (https://github.com/schatzlab/pseudohaploid) After each of these correction steps, completeness of the assembly was assessed by identifying Benchmarking Universal Single-Copy Orthologs (BUSCOs) using the BUSCO (v3.0.2) pipeline in genome mode (51). We identified single-copy orthologs based on the insecta– db9. Each of the correction steps improved the assembly quality, especially with regard to BUSCO completeness scores (Table 2).

**Table 2.**
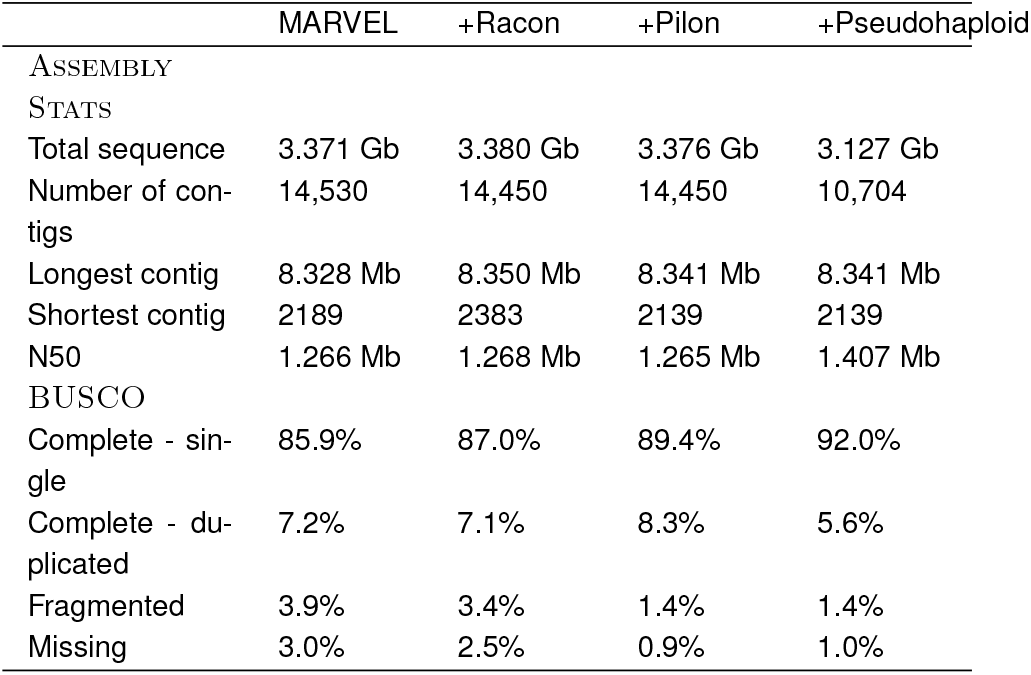
Assembly statistics and BUSCO scores for each assembly stage.

### Repeat annotation

Repetitive elements from *D. punctata* genome assmebly were categorised with repeat modeler (http://www.repeatmasker.org/), LTRharvest (52) and TransposonPSI (http://transposonpsi.sourceforge.net). The resulting libraries were merged together with the SINEbank repeat data base, specific to Insecta (53). The merged repeat library was filtered for redundancy using cd-hit-est (parameters: -c 0.8 -n 5) (54) and for true proteins by blasting against a *de novo* assembled *Diploptera punctata* transcriptome. Specifically, we generated the *de novo* transcriptome assembly with 29 previously published RNAseq libraries (6) using Trinity (55) at default settings. Nucleotide coding and protein sequences were generated from the Trinity assembly with TransDecoder (http://transdecoder.github.io/). Sequences were removed from this transcriptome if they received a significant blast hit (e-value < 1e-5) against the RepeatPeps library contained in the RepeatMasker data set. We then blasted our merged repeat library against this reduced set of transcripts using blastn. Any hits with an e-value < 1e-10 were removed from the library. The repeat library was classified with RepeatClassifier. The genome assembly was then soft masked with RepeatMasker.

### Gene annotation

We used two programs to predict *ab initio* gene models: Braker (56), which combines Augustus (57) and GeneMark (58), and GeMoMa (59). Both were trained with the *Blattella germanica* genome and *D. punctata* RNAseq (6). We additionally used two methods of evidence-based gene prediction. With Spaln (v2.4.6) (60) we aligned a large database of proteins against our genome assembly. The protein database contained the Uniprot arthropod database (version April 2018) and all available Blattodea proteomes: *B. germanica* (16), *Periplaneta americana*, *Cryptotermus secundus* (16), *Zootermopsis nevadensis* (61) and *Macrotermes natalensis* (62). Finally gene models with predicted by aligning the RNAseq data to the genome assembly with Pasa (63). EVidenceModeler (64) was then used to combine the different gene sets. The following weights were applied to each gene set; Augustus and GeneMark: 1; GeMoMa: 2; Spaln: 5; Pasa: 10. This produced a GFF containing 61,692 putative protein coding genes, which was further filtered to remove contamination and repetitive elements using blast against the NCBI nr database and our repeat database, respectively. Annotation scores from EVM output were compared to noncoding equivalent. All putative genes with an annotation score =< noncoding equivalent were removed. Furthermore, to detect true positives, PFAM domains were annotated on translated sequences with pfamscan and RNAseq reads were mapped against the putative gene regions. All gene models with at least one significant PFAM domain or to which at least 10 reads mapped in at least one sample were considered true positives and retained. All further genes were only kept if supported by evidence from protein alignments (Spaln), transcript alignments (Pasa), or homology within Metazoa, resulting in a gene set of 27,940 protein coding genes.

Chemoreceptor genes are notoriously difficult to predict with standard tools and were therefore annotated manually with bitacora (65) and exonerate (66) in two rounds. For the first round, the chemoreceptors from *Blattella germanica* (25), *Drosophila melanogaster*, *Apis melifera* and *Apolygus lucorum* species were taken as a database for bitacora and exonerate. Predicted gene models were filtered for the presence of domains of interest and length (85% of domain length average) and used as a database for the second round. The filtered predictions were merged between bitacora and exonerate and with the previous annotations. Predicted cuticle proteins were identified with BLAST comparison to known protein in other insect system (26).

### Ortholog detection and phylogeny

The 18 Insect species have been carefully chosen to investigate viviparous transition as it encompasses the independent origins of viviparity in different insect orders along with, at least, 1 outgroup species per order. Orthologs among the insect species selected; including mayfly (*Ephemera danica*) used as a general insect outgroup, whitefly (*Bemisia tabaci*) as outgroup for the Hemiptera branch, 2 species of aphids (*Acyrthosiphon pisum*, *Rhopalosiphum maidis*) representing origin of viviparity in Hemiptera, 4 species of Diptera (*Glossina morsitans* as viviparous dipteran, and *Musca domesticus*, *Drosophila melanogaster*, *Stomoxys calcitrans* as dipteran outgroups), 2 species of stick insects (*Medauroidea extradentata*, *Clitarchus hookeri*) and locust (*Locusta migratoria*) as outgroup of Blattodea, 3 species of cockroaches (*Blattella germanica* and *Periplaneta americana* as outgroups, *Diploptera punctata* as viviparous blattodean), and 4 species of termites (*Zootermopsis nevadensis*, *Cryptotermes secundus*, *Coptotermes formosanus*, *Macrotermes natalensis*); were discovered using Orthofinder (v2.5.2) (18). To optimize the number of singlecopy orthologs, We categorized them as such if they were single-copy or absent in viviparous species and oviparous species with multiple copies were dropped. Ortholog families including at least one viviparous species and three oviparous species were retained for further analyses, culminating at 5463 single-copy ortholog families. Phylogenetic tree reconstruction, including all species described above, was undertaken by OrthoFinder (18).

### Duplication and loss events

The variation of gene family size across the phylogenetic tree was assessed with CAFE (v4.2.1) (19), to unravel the expansion and contraction of gene families in all branches of the tree. Moreover, the variation of domain numbers in across the phylogenetic tree was assessed with CAFE (v4.2.1) (19).

### Multiple Alignment

For each single-copy ortholog family, the longest protein isoforms for each of the species gene were used in multiple sequence alignment with PRANK (v.150803) (67) and unreliably aligned residues and sequences were masked with GUIDANCE (v2.02) (68). This combination was shown to perform the best on simulated data (69). To optimize alignment length without gaps, we ran maxalign script (70) and removed subsequent sequences leading to more than 30% of gapped alignment as long as it did not result in the removal of a viviparous species’ sequence, and an alignment of less than 4 sequences. The protein sequences were replaced with coding sequences in the multiple alignments using pal2nal script (71). Alignments regions, where gapped positions were present, were removed with a custom python script (see Supplementary Code SC1), as these are the most problematic for positive selection inference (72). Finally, CDS shorter than 100 nucleotides were eliminated (73). After filtering, our dataset included 4,671 gene families. The mean length of filtered alignment was 614 nucleotides (median = 471 nucleotides), ranging from a minimum of 102 nucleotides to a maximum of 7836 nucleotides and included on average 10 sequences (median = 11), ranging from 4 to 18.

### Identifying selection pressures

#### Branch-Site tests to detect positive selection

Phylogenetic tests of positive selection in protein-coding genes usually contrast substitution rates at non-synonymous sites to substitution rates at synonymous sites taken as a proxy to neutral rates of evolution. The adaptive branch-site random effects model (aBSREL, (28)) from Hyphy software package (27) was used to detect positive selection experienced by a gene family in a subset of sites in a specific branch of its phylogenetic tree. Test for positive selection was run only on the branches leading to the origin of viviparity, namely the *Diploptera punctata* branch, the *Glossina morsitans* branch, and the aphid branch. Results from the adaptive branch-site random effects model were corrected for multiple testing as one series using False Discovery Rate (FDR) (74) and set up our significant threshold at 10% (31).

#### Variation of Selection pressure

While elevated dN/dS can be caused by increased positive selection, it can also be the result of relaxed purifying selection, or a combination of both. We used RELAX (32) to categorize if shifts in the distribution of dN/dS across individual genes are caused by overall relaxation of selection (i.e. weakening of both purifying selection and positive selection, towards neutrality) versus overall intensification of selection (i.e. strengthening of both purifying selection and positive selection, away from neutrality). Specifically, RELAX models the distribution of three categories of dN/dS (i.e. positive selection, neutral evolution, purifying selection) across a phylogeny and compares the distributions for foreground branches (here, the branches of viviparous origins) to background branches (here, the ancestral and sister branches of viviparous origins) and estimates a parameter K that indicates overall relaxation (*K* < 1) or intensification (*K >* 1). Eight alignments failed to run due to removal of reference species during filtering. Results from RELAX models were corrected for multiple testing as one series using (FDR) (74) and set up our significant threshold at 10% (31).

### Test for functional category enrichment

Gene Ontology (GO) (75) annotations for our gene families were taken from pfam annotations and from orthologs of *Drosophila melanogaster* and the enrichment of functional categories was evaluated with the package topGO version 2.4 (20) of Bioconductor (76).

To identify functional categories enriched for expanded and contracted gene families, the Fisher exact test with the ‘elim’ algorithm of topGO was run separately for the significantly expanded and contracted gene families which were given the score of 1 while other gene families were given the score of 0. The results were then corrected with the FDR to account for multiple testing (74) and set up our significant threshold at 20% (31). Gene Ontology categories mapped to less than 10 genes were discarded.

To identify functional categories enriched for genes under positive selection, strengthened, and relaxed selection pressure, the SUMSTAT test was used as described in (31). The SUMSTAT test is more sensitive than other methods, and minimizes the rate of false positives (77). To be able to use the distribution of loglikelihood ratios of the aBSREL and RELAX tests as scores in the SUMSTAT test, a fourth root transformation was used (31). This transformation conserves the ranks of gene families (78). In addition, we assigned a log-likelihood ratios of zero for genes under relaxed selection (*K* < 1) when testing for enrichment of functional categories with genes under strengthened selection and vice-versa (0 for genes with *K >* 1) when testing for enrichment from genes under relaxed selection. Gene Ontology categories mapped to less than 10 genes were discarded.

The list of significant gene sets resulting from enrichment tests is usually highly redundant. We therefore implemented the “elim” algorithm from the Bioconductor package topGO, to decorrelate the graph structure of the Gene Ontology (20). To account for multiple testing, the final list of p-values resulting from this test was corrected with the FDR (74) and set up our significant threshold at 20% (31). To cluster the long list of significant functional categories before FDR correction, we used REVIGO (30) with the SimRel semantic similarity algorithm and medium size (0.7) result list.

### RNA-seq analyses

To assess if transcriptional changes are similar between viviparous insects, previously available RNA-seq data sets of multiple *Glossina* species and *D. punctata* were analyzed (6, 26). In specific, this allowed comparison between males, non-pregnant females, early pregnancy, and late pregnancy. Transcripts per million (TPM) was determined using Sailfish (79). The expressional changes were compared with the weighted correlation network analysis (WGCNA). In specific, orthologs that were identified through the use of Orthofinder (18) between *Diploptera* and *Glossina* sp. with sequenced genomes (6, 26). The single copy orthologs obtained from Orthofinder were used for WGCNA to identify groups of genes with similar expression profiles males, during early pregnancy, and late pregnancy. The WGCNA was conducted as a signed analyses with a soft power of 12. Modules that were significantly associated with early and late pregnancy were analyzed for enriched GOs following a false detection rate detection. Immune-related genes were analyzed through the use of Deseq of previous data (6), where the putative immune genes were identified. Clustering was performed on normalised counts using the R package DEGreports v. 1.25.1 (80), with a minimum cluster size of 20.

### Functional validation of factors identified in genomics and transcriptomic studies

Together with our results of detection of selection in viviparous species and gene’s expression at different life-stages in *D. punctata*, we identified genes of interest, which could be linked with the adaptation to viviparity. RNA interference was conducted according to (6) and (81). Briefly, dsRNA was generated with a MEGAscript RNAi Kit (Ambion). Following preparation of the dsRNA, each pregnancy female was injected with 2-3 μg of dsRNA at 30-40 days into the pregnancy cycle with a pulled glass capillary needle. Control individuals were injected with a dsRNA targeting green fluorescent protein (6). Individuals were monitored for abortions and the duration of pregnancy.

Immune functionality was assessed through the use of injection of *Pseudomonas aeruginosa* in pregnant females to confirm potential altered immunity during this state. To do so, bacteria was grown until a log-phase and injected 1.0 × 10^5^ CFU of *P. aeruginosa* in 3 l PBS in pregnant or virgin females. Survival was monitored for 40 days.

## ACKNOWLEDGMENTS

We thank David L. Denlinger for comments and suggestions on an early draft of this manuscript and Carsten Kemena, Elias Dohmen, and Anna Grandchamp for their help in troubleshooting the comparative genomics pipeline. This work was supported by a Discovery grant, no. 9408-08 from the Natural Sciences and Engineering Research Council of Canada to S.S.T, National Institute of Allergy and Infectious Diseases of the National Institutes of Health under Award Number R01AI148551 and National Science Foundation DEB1654417 (to J.B.B. for shared computational resources), B.F. was supported by an EU-H2020 MSCA-IF-2020 fellowship (101024100, TEEPI), M.C.H. was supported by a DFG grant BO2544/11-2 to E.B.B, A.M. was supported by a DFG grant HA8997/1-1 to M.C.H. and S.E. was supported by a Royal Society Dorothy Hodgkin Fellowship (DH140236).

